# Identifying cortical areas that underlie the transformation from retinal to world motion signals

**DOI:** 10.1101/2022.06.14.496150

**Authors:** Puti Wen, Michael S. Landy, Bas Rokers

**Affiliations:** Psychology, New York University Abu Dhabi; Dept. of Psychology and Center for Neural Science, New York University; Psychology, New York University Abu Dhabi, Dept. of Psychology and Center for Neural Science, New York University

## Abstract

Accurate motion perception requires that the visual system integrate the retinal motion signals received by the two eyes into a single representation of 3D (i.e., world) motion. However, most experimental paradigms limit the motion stimuli to the fronto-parallel plane (i.e., 2D motion) and are thus unable to dissociate retinal and world motion signals. Here, we used stereoscopic displays to present separate motion signals to the two eyes and examined their representation in visual cortex using fMRI. Specifically, we presented random-dot motion stimuli that produced percepts of various 3D motion trajectories. We also presented control stimuli that contained the same retinal motion energy in the two eyes but were inconsistent with any 3D motion trajectory. We decoded the stimuli from BOLD activity using a probabilistic decoding algorithm. We found that 3D motion direction can be reliably decoded in three major clusters in the human visual system. In early visual cortex (V1-V3), we found no significant difference in decoding performance between the 3D motion and control stimuli, suggesting that these areas represent retinal motion rather than world motion signals. In voxels in and surrounding hMT and IPS0 however, decoding performance was consistently superior for 3D motion compared to control stimuli. Our results reveal the parts of the visual processing hierarchy that are critical for the transformation of retinal into world motion signals and suggest a role for IPS0 in the representation of 3D motion signals, in addition to its sensitivity to 3D object structure and static depth.

## Introduction

From escaping predators to hunting prey, the ability to accurately perceive motion has enabled our survival for as long as we have existed. Although most humans may no longer need to run away from bears, daily activities, such as pouring coffee, driving, and playing sports, are still critically dependent on motion perception. In squash, a player has moments to decide when and where to swing. To support this simple action, the brain carries out tricky perceptual tasks such as motion detection (Borst and Egelhaaf 1989), the judgment of direction (Britten et al. 1992; Salzman, Britten, and Newsome 1990), speed (Liu and Newsome 2005), and time-to-contact (Tresilian 1995) in the blink of an eye.

At the cortical level, the visual system processes these motion signals along a hierarchical pathway (Maunsell and van Essen 1983; Van Essen et al. 1986; Maunsell 1992; DeYoe et al. 1994). Primary visual cortex (V1) and the middle temporal area (MT) are two key sites in this pathway, where the former is thought to primarily process component motion signals and the latter pattern motion signals (Britten et al. 1993; Movshon and Newsome 1996; Born and Bradley 2005; Rust et al. 2006). However, it is not yet clear how the representation of motion contained in the 2D retinal images is transformed into a representation of motion trajectories in three dimensions, or where this transformation takes place.

Multiple studies have investigated neural responses to stimuli that either imply or contain motion-in-depth cues. Monkey MT neurons are organized in columns by both motion direction (Albright, Desimone, and Gross 1984) and disparity preference (DeAngelis and Newsome 1999). About half of neurons in macaque area MT are selective to motion-in-depth direction (Sanada and DeAngelis 2014; Czuba et al. 2014). Moreover, MT has been linked to the perception of 3D structure from motion (Bradley, Chang, and Andersen 1998; Grunewald, Bradley, and Andersen 2002), and depth from both binocular disparity and binocular motion (Nadler, Angelaki, and DeAngelis 2008; Armendariz et al. 2019). Human fMRI studies have provided evidence of adaptation and motion aftereffects in hMT+ to 3D motion direction (Rokers, Cormack, and Huk 2009; Czuba et al. 2011; Joo et al. 2016).

Recent work in macaque monkeys has found that decoding of 3D motion in MT is best explained by the monocular motion signals of the dominant eye and suggested that, at least in non-human primates, 3D motion processing may happen in additional regions such as area FST (Héjja-Brichard et al. 2020). Similar arguments have been made based on results in humans (Likova and Tyler 2007). Furthermore, research on depth perception (Ban et al. 2012) and structure-from-motion (Orban et al. 1999; Georgieva et al. 2009) has suggested that areas such as V3A/B and IPS may play a role in the extraction of 3D motion signals as well.

Here, we used an fMRI-based decoding approach to understand the representation of binocular stimuli that signal 3D motion across the visual hierarchy. We presented various trajectories of 3D motion to observers and asked whether and where we can decode 3D motion direction from the BOLD response. To test that a given cortical area truly represents 3D motion direction rather than aspects of the two monocular retinal motion signals alone, we analyzed responses to a control stimulus in which the retinal signals moved vertically, rather than horizontally. This control stimulus matched the motion energy in the horizontal motion stimulus, but because of the geometry of binocular motion perception, did not produce 3D motion percepts. Specifically, because of the horizontal offset of our two eyes, horizontally opposite retinal motion is consistent with a stimulus approaching or receding from the observer. Vertically opposite retinal motion on the other hand is not consistent with any given 3D motion trajectory. Moreover, since we used relatively sparse dot motion displays, the stimuli did not induce binocular rivalry, but instead produced percepts of transparent 2D motion (see Supplementary Videos 1 and 2). We hypothesized decoding performance for 3D-motion and control stimuli would be equivalent in visual areas that encode retinal motion. However, cortical regions that integrate the retinal motion signals from the two eyes, should exhibit worse decoding performance for the vertical compared to the horizontal motion stimuli.

In summary, we aimed to elaborate our understanding of the classical motion processing pathway using binocular stimuli that can signal 3D motion. As 3D motion processing relies on neural mechanisms beyond those required for the processing of fronto-parallel motion, we expected that a critical evaluation of the BOLD responses produced by our 3D motion stimuli would reveal a more complete picture of the visual motion processing hierarchy in the human brain.

## Materials and Methods

### Observers

Nine healthy observers (one male, age 23-46 years) with normal or corrected-to-normal vision participated and provided written informed consent. All observers achieved a score of five or higher (70 seconds of arc or better) on the Randot Circles Stereotest (Stereo Optical Company, Chicago, IL). Four observers participated at New York University Abu Dhabi, and five observers participated at New York University New York. All observers participated in two scanning sessions for the horizontal condition and two sessions for the vertical condition. Each scanning session lasted 1.5 hours. The experiment was approved by the University Committee on Activities Involving Human Subjects at New York University and New York University Abu Dhabi.

### Visual Stimulus

Stimuli were generated on a Macintosh computer using MATLAB 9.2 (The MathWorks, Natick, MA, USA) and the Psychophysics Toolbox extensions (Brainard 1997; Pelli 1997; Kleiner et al. 2007). We presented the binocular stimuli using a ProPixx DLP LED projector (VPixx Technologies Inc., Saint-Bruno-de-Montarville, QC, Canada; screen resolution: 1920 × 1080 pixels, refresh rate: 120 Hz; 60 Hz for each eye) with a rear projection screen (projected screen width: 38.5 cm, viewing distance: 88 cm) positioned at the back of the scanner.

The stimuli were white dots (55.2 cd/m^2^) with 0.5 s lifetime drifting against a black (0.2 cd/m^2^) background (**Fig. 1A**^**1**^). All dots were restricted to two black sectors in the left or right hemifield (outer aperture radius: 14 deg, inner aperture radius: 0.7 deg). No dots were presented in two wedges centered on the upper and lower vertical meridians (25 deg of arc wedge width). The wedges allowed us to present stimuli with opposite motion directions on either wedge side, reducing oculo-motor drive (see below). We ensured that dots in the two eyes remained within the sector and remained binocularly visible at all times. The remainder of the display area was filled with 1/*f* noise that was identical in the two eyes and perceptually appeared at the fixation distance. There was a fixation dot at the center of the display.

**Fig. 1.**
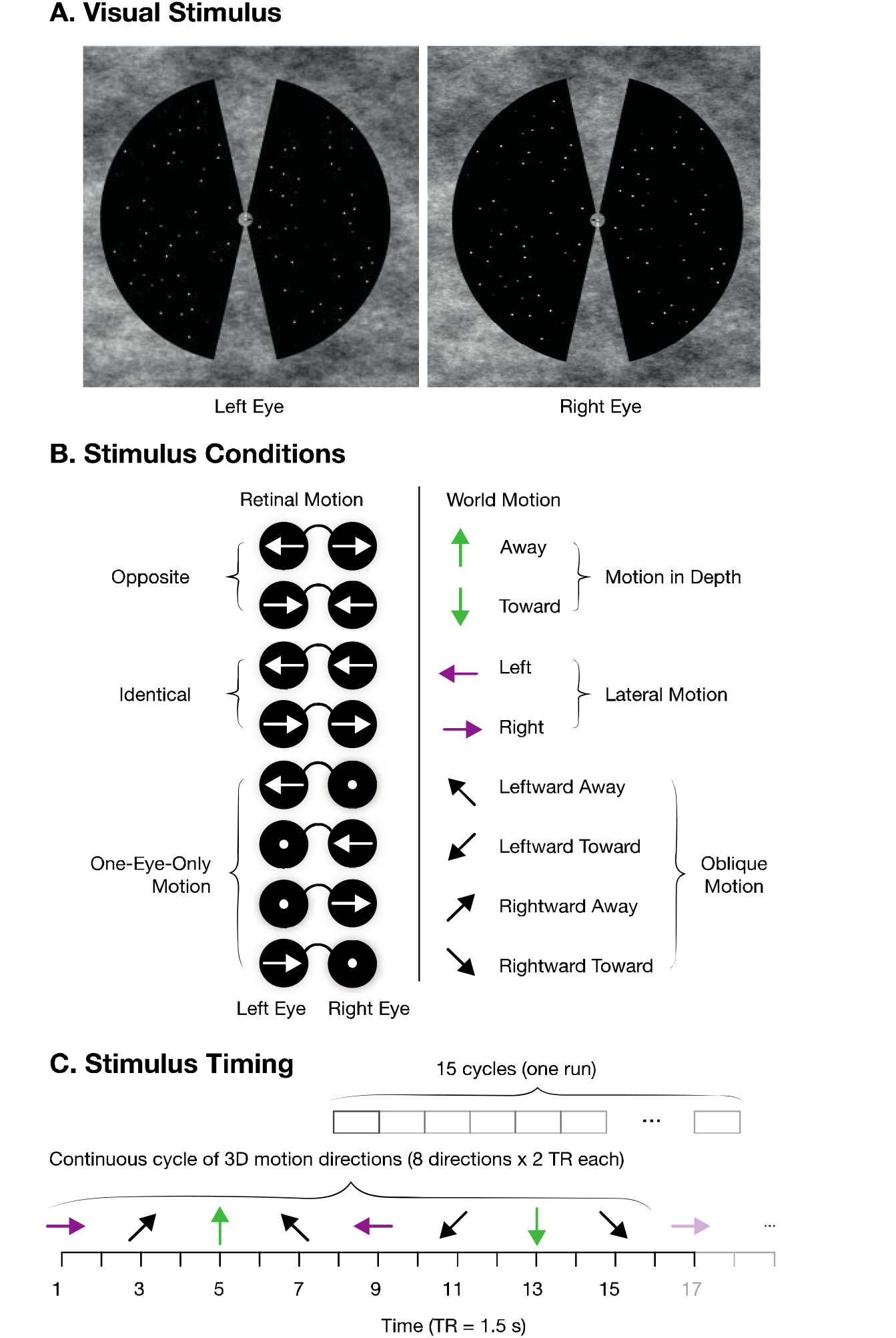
Stimuli and experimental design for the main experiment (horizontal condition). ***A. Visual Stimulus***. In each monocular image, white dots drifted horizontally against a black background. Dots were presented within a circular aperture filled with pink noise to aid fixation and binocular fusion. At all times, opposite motion directions were presented in the two hemifields. ***B. Stimulus Conditions***. The retinal motion signals produce eight different world-motion directions, depending on the relationship of retinal motion between the two eyes. ***C. Stimulus Timing***. Timeline for one run of the main experiment. There were 15 cycles of each of the eight directions after removing the first ten volumes of the scan. In the odd runs, the stimuli appear in a continuous counterclockwise cycle of 3D motion directions. The order was flipped to a clockwise cycle in the even runs.

The total number of white dots displayed in both hemifields at a time for each eye remained at 100 as a new dot would appear at a random location on the screen when a previous dot disappeared. To reduce false binocular matches, any new dot position was set to a minimum of 0.975 deg absolute Euclidean distance from the rest of the dots that were currently on the screen. We restricted the maximum horizontal dot disparity between the two eyes to 0.3 deg. At the onset of the experiment, each dot was initialized with a random disparity chosen from a uniform distribution between the minimum (−0.3 deg) and maximum (0.3 deg) disparity. Depending on the relative velocity of the dots presented to the left and right eye, the observers perceived different motion directions. Dots drifted horizontally in the horizonal condition and vertically in the vertical condition. All other stimulus parameters were identical between the two conditions.

### Horizontally opposite motion stimuli

The main experiment contained three types of motion direction: lateral, in-depth, or oblique (**Fig. 1B**). Each dot in the lateral-motion stimuli maintained a constant disparity throughout its lifetime, moving strictly leftward or rightward at a rate of 0.85 deg/sec. Dots in the motion-in-depth stimuli changed disparity at 1.7 deg/sec (dot speeds of 0.85 deg/sec in each eye but in opposite directions), with no net leftward or rightward motion. Oblique motion stimuli were a combination of lateral and in-depth motion: dots moved leftward or rightward at a rate of 1.2 deg/sec in one eye and were stationary in the other eye. These dot speeds were used because they allowed for high direction-discrimination sensitivity for 3D motion (Czuba et al. 2010).

Stimuli cycled through motion directions either clockwise or counterclockwise in the horizontal (xz) plane. For example, when cycling counterclockwise rightward motion was followed by right/away, away, left/away, left, left/toward, toward, right/toward, right again and so on (**Fig. 1C**). The eight possible lateral/in-depth/oblique motion directions were presented for three seconds each.

Each MRI scan run lasted 375 seconds in total. The first 15 seconds (5 trials) of each run were removed to minimize the effects of transient magnetic saturation and to allow for the hemodynamic response to reach a steady state. In the remaining 360 seconds, each cycle of eight directions was repeated 15 times. Direction order was alternated (clockwise vs. anti-clockwise) from scan to scan. At all times, opposite motion directions were presented in the two hemifields. For example, if dots in the right hemifield were moving away from the observer, then dots in the left hemifield would move towards the observer. This was done to discourage eye movements induced by the optokinetic response and encourage consistent fixation on the center of the screen.

### Vertically opposite motion stimuli

All participants also took part in a vertical-motion experiment. All stimulus properties were identical to that in the main experiment, with the exception that dots drifted vertically rather than horizontally **(Fig. 2**^**2**^ ). Thus, trial timing as well as stimulus parameters such as speed and luminance were maintained between experiments, with the critical difference that the vertically opposite motion stimuli produce transparent motion, rather than 3D motion percepts.

**Fig. 2.**
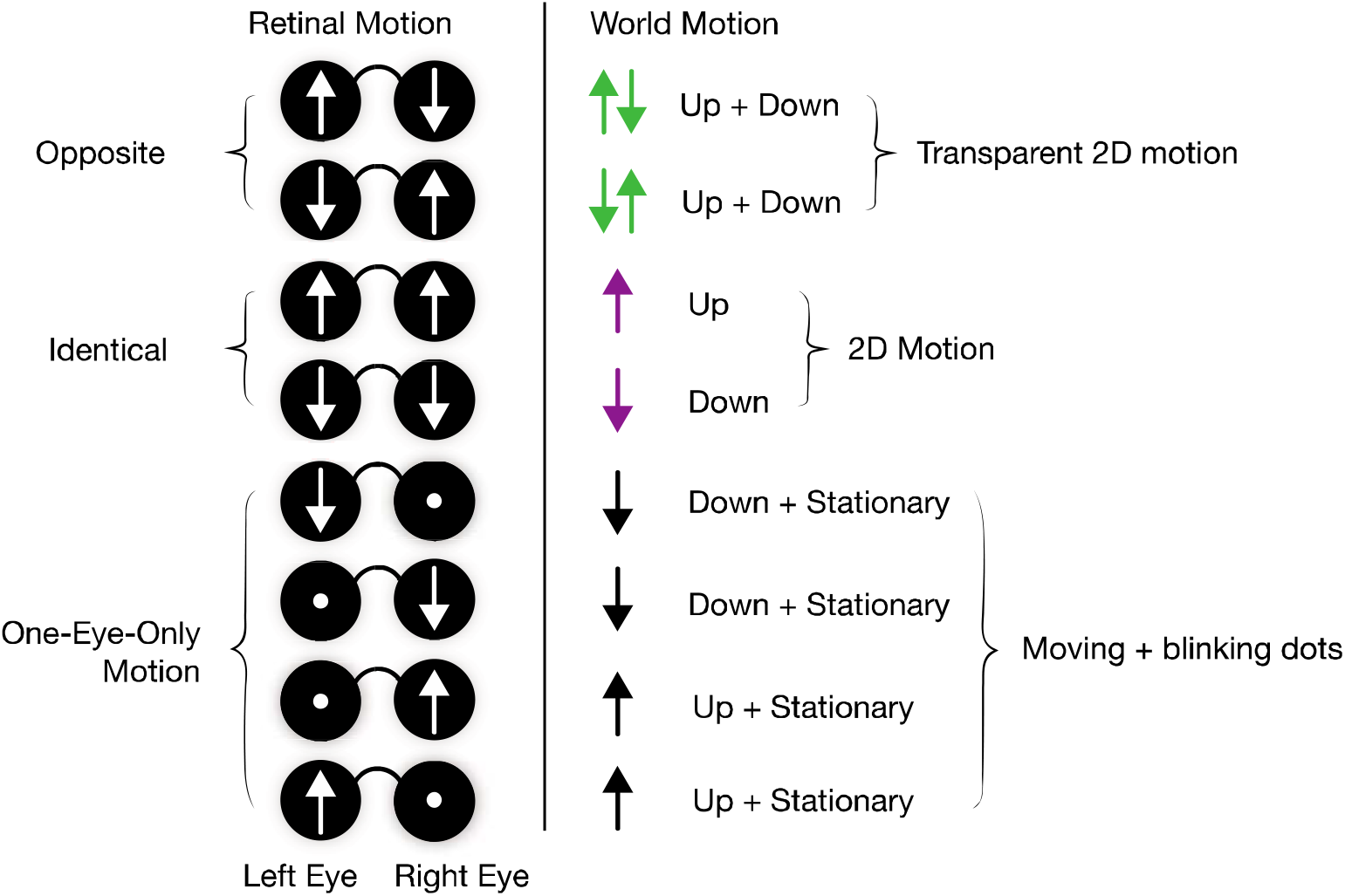
Stimuli and experimental design for the vertical condition. Each horizontal retinal motion trajectory was rotated by 90 deg and now moved vertically. The control experiment has analogous retinal stimulus properties while no longer producing 3D motion percepts. Opposite retinal motion (upward motion in one eye and downward motion in the other) produces a percept of transparent motion rather than of motion in depth.

### Behavioral task

To control attention and ensure proper vergence throughout the experiment, subjects performed a depth-detection task at fixation. At random intervals, the binocular disparity of the fixation dot changed slightly (.1 deg of arc or less). Participants indicated the perceived depth change (near/far disparity relative to the screen) by button press.

### MRI data acquisition

MRI data were acquired on a Siemens Prisma 3T full-body MRI scanner (Siemens Medical Solutions, Erlangen, Germany) using a 64-channel head coil. For each observer, a T1-weighted anatomical scan was acquired (TR: 2400 ms; TE: 2.22; flip angle: 8°; 0.8 mm isotropic voxels). This anatomical volume was used for white/gray matter segmentation and co-registration with the functional scans. T2*-weighted functional scans were acquired using an echo-planar imaging (EPI) sequence (TR: 1500 ms; TE: 38 ms; flip angle: 68°, multiband factor: 4; matrix size: 104 × 104, 2 mm isotropic voxels; 68 slices). For each functional scan, we collected 250 volumes with full brain coverage.

### Anatomy and ROI Identification

We anatomically identified visual cortical ROIs using the Neuropythy atlas (Benson and Winawer 2018). The following ROIs were selected and used in the analysis: V1v, V1d, V2v, V2d, V3v, V3d, V3A, V3B, hV4, VO1, VO2, PHC1, PHC2, IPS0-5, hMT, MST, LO1, LO2, SPL1, and FEF. We merged V1-3v and V1-3d as V1-3 using *fslmaths*.

### fMRI Pre-processing

All scans were organized in the Brain Imaging Data Structure (BIDS) format (Gorgolewski et al. 2017) using dicom2bids and dcm2niix (Li et al. 2016) and then pre-processed using the fMRIPrep pipeline (version 20.2.1) for motion correction, spatial normalization, and co-registration between the functional and anatomical scans (Esteban et al. 2019).

### Decoding motion directions

For functional runs and for each ROI, the data were organized as an *n* X *m* matrix where *n* is the number of measurements (*n* = 240) for each scan and *m* is the number of voxels within that ROI. Every column in this matrix was a time series of raw fMRI signals for one voxel. Prior to classification, we applied a high-pass filter using the fast Fourier transform (FFT) to convert the data from the time domain to the frequency domain and removed low-frequency noise (cutoff: 1/72 Hz). The remaining data were then converted back to the time domain with the inverse FFT. We then normalized (*z*-scored) the time series separately for each voxel. Each scan contained 120 trials – 15 trials for each of the eight motion directions – and each trial lasted 2 TRs (3s). We averaged the trials within a scan that had the same motion direction 6 s after the stimulus onset to account for the hemodynamic delay. This resulted in an eight (number of directions) by *m* (number of voxels) matrix for each scan. Each row of the final matrix represents the average normalized BOLD response to a given motion direction, and each column represents the data for one given voxel. Pre-processing was performed for each subject, and each scan separately.

We decoded the motion direction using a probabilistic decoding algorithm, TAFKAP (van Bergen and Jehee 2021). This is an inverted generative model that decodes both a stimulus estimate and its uncertainty. Since we only used eight motion directions, we used an extension of TAFKAP to discrete stimulus sets (van Bergen, pers. comm.). This algorithm returns the direction with the highest probability as the estimated direction and computes the entropy of the decoded distribution as its measure of uncertainty. We applied 32-fold cross-validation and saved the averaged decoding results across validation in an eight-by-eight confusion matrix, where each column represents a presented direction, and each row records the proportion of scans in which the direction was decoded as each of the eight possible directions. Accurate decoding was reflected on the diagonal of this matrix. We obtained a bootstrapped distribution of decoding results for chance performance based on repeatedly randomizing direction labels for the training data. We considered the decoding accuracy for the original data to be statistically significant if the value exceeded the 95th percentile of this chance-performance distribution.

We used the MATLAB function *classify* with the option ‘diaglinear’ to decode motion directions in the searchlight analysis. The classifier fit a multivariate normal density to each motion direction with a diagonal covariance matrix using the training data and its label. The diagonal covariance matrix represented an assumption of independent responses across voxels since the data were not sufficient for an accurate estimate of the true covariance. We used the 80:20 training-to-testing-split method to cross-validate the classifier. We resampled data with replacement 5000 times to compute the chance distribution with randomized labels. The *p*-value was calculated as the percentage of times the resampled chance decoding accuracy was larger than the observed decoding accuracy.

### Motion direction preferences

We converted the data into fsaverage6 surface space using FreeSurfer (Reuter et al. 2012). We computed the preferred motion direction for each vertex. We first averaged the BOLD response across trials and scans for each presented motion direction. We then computed the vector sum across motion directions, so that the vector length represents the averaged BOLD response, and the angle represents the preferred motion direction of that vertex. The vector length can also be understood as the strength of the motion direction preference. We computed this vector for each vertex and each subject separately and then calculated the circular mean of this vector across subjects for each vertex. We define the mean vector as the preferred motion direction for that vertex. We plotted the result on a flattened surface map for both left and right hemispheres and thresholded the figure by motion preference strength (top 10th percentile).

## Results

### Reliable decoding of 3D motion in V1 and hMT

We probed the representation of 3D motion signals in the canonical motion-sensitive areas, V1 and MT. We presented separate motion stimuli to the two eyes, consistent with a 3D volume of dots that changed their direction every 3 seconds, cycling through 8 motion directions every 24 seconds. We extracted the mean response in every voxel to each motion direction in each of 20 scans, and trained a classifier to decode the presented motion direction in each ROI. The average decoding results (N=9) are illustrated in **Fig. 3A**. Correct classification, where the decoded direction matched the presented direction, corresponds to elements on the positive matrix diagonal. Average decoding performance across all directions was 41.19 % (+/- 3.88% SE) and 35.2 % (+/- 2.32% SE) in V1 and hMT respectively, reflected in a prominent red positive diagonal across the confusion matrices in both V1 and hMT.

**Figure 3.**
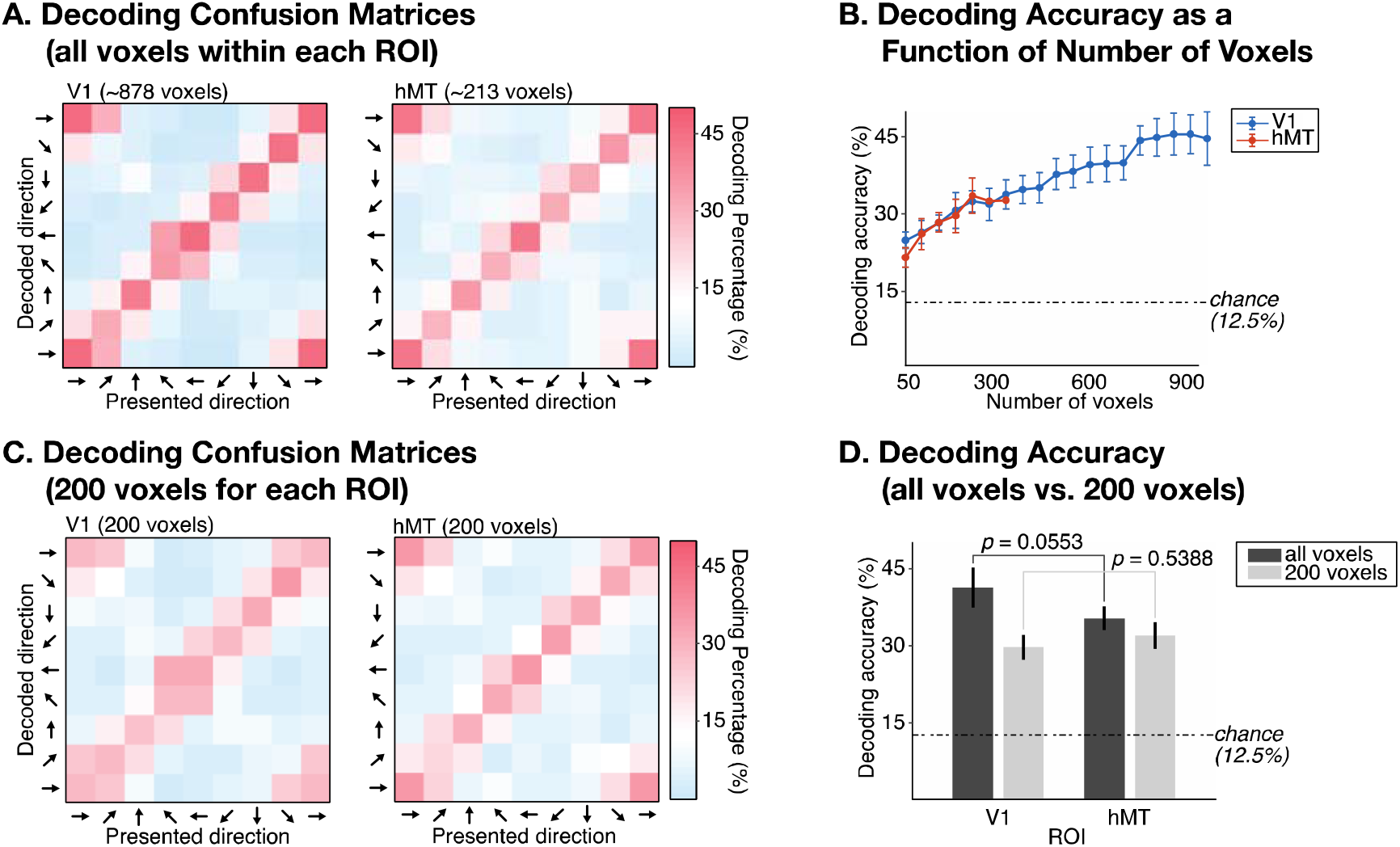
Reliable decoding of 3D motion signals in both V1 and hMT. ***A***, Decoding confusion matrices using all voxels in V1 and hMT. Matrices are color-coded based on decoding percentage. Theoretical chance performance (12.5%) in white, above chance performance in red, and below chance in blue. The first row and column of the matrix are repeated at the end of each matrix to illustrate the circular nature of the eight motion directions. ***B***, Decoding accuracy as a function of the number of voxels included in V1 and hMT. Error bars indicate SEM across subjects. Dashed black lines indicate the theoretical chance of decoding one of eight motion directions (12.5%). ***C***, Same analysis as ***A*** except only 200 randomly-selected voxels from each ROI were used. ***D***, Decoding accuracy between using all voxels and using only 200 voxels for V1 and hMT. Error bars indicate SEM across subjects. Dashed black lines indicate the theoretical chance of decoding one of eight motion directions (12.5%). All data reflects results averaged across participants (*n* = 9).

### Accounting for the number of voxels

A common finding in the decoding of motion direction from BOLD responses, is that decoding accuracy is greater in V1 than MT (Kamitani and Tong 2006; Serences and Boynton 2007; Wang et al. 2014), even when equating for the difference in size between these areas. For example, at our 2 mm isotropic imaging resolution, V1 contained ∼878 voxels, and hMT only contained ∼213 voxels on average (*n* = 9). To equate for area size, we re-ran the classifier in V1 and hMT, each time increasing by 50 the number of voxels used for decoding. When equating the number of voxels, decoding accuracy was near-identical in V1 and hMT (**Fig. 3B**). Thus, in contrast to prior work on the decoding of 2D motion, we found no evidence for a difference in decoding accuracy between V1 and MT.

Prior work has suggested that superior performance in the decoding of motion (and orientation) in early cortical visual areas, including V1, may be due to vignetting, biases in the BOLD response to visual stimuli near the edges of stimulus apertures (Carlson 2014; Wang et al. 2014). Our results suggest that these biases may be less pronounced for stimuli that are not constrained to the image plane, a question we will return to at the end of the Results section.

To directly compare decoding performance across ROIs, we set 200 voxels as the upper limit for every ROI for all subsequent analyses. If an ROI had more than 200 voxels, we randomly selected 200 voxels for each cross-validation. Limiting the number of voxels did impact classifier performance, reducing decoding performance in both V1 (*t*_*V1*_(8) = 4.74, *p* = .002) and hMT (*t*_*hMT*_(8) = 3.49, *p* = .008), although the reduction in the percentage of accurately decoded stimuli in V1 (from 41.2% to 29.6%) was significantly larger than in hMT (from 35.2% to 31.8%; *t*_*V1-hMT*_(8) = 3.41, *p* = .009). However, the overall pattern of results remained relatively unaffected (compare **Figs. 3A** and **Fig. 3C**), where a red positive diagonal line is still apparent in both regions, although now less prominent in V1. In contrast to all previous work (Kamitani and Tong 2006; Serences and Boynton 2007; Wang et al. 2014), decoding in hMT is no worse than in V1 once equating for number of voxels (*t*_*V1-hMT(200)*_(8) = -.64, *p* = .539; **Fig. 3D**). Having put overall decoding performance in V1 and MT on equal footing, we next turned to understanding differences in the representation of motion signals in each region, based on two additional analyses.

### Decoding misclassification

Reliable and comparable decoding of 3D motion signals in V1 and MT does not necessarily imply classification based on similar information. Inspection of the confusion matrices indicates that decoding errors were not randomly distributed. In the confusion matrices, the off-diagonal squares represent the percentage of trials the presented motion direction is misclassified based on the pattern of BOLD responses across voxels in an ROI. The decoder often misclassifies the presented motion direction as an immediately neighboring one. For example, the decoder will often misclassify rightward-away oblique motion as either rightward or away rather than any other motion direction. Conversely, the decoder rarely misclassifies the presented motion direction with a direction 180 degrees away (e.g., misclassifying leftward as rightward or approaching as receding motion). As a result, many red diagonal lines appear to be three squares wide and are bordered by blue regions with below-chance decoding (**Fig. 3C**).

The oblique motion direction stimuli in our experimental design only contain motion signals in one eye. For example, the rightward-away motion stimulus presents rightward motion to the right eye, but a stationary field of dots to the left eye **(Fig. 1B)**. If a voxel in an ROI predominantly encodes (monocular) retinal input rather than world motion, we expect that ROI to show a bias towards the presented monocular retinal motion. As monocular visual inputs integrate and transform into a representation of world motion, we expect a decrease in that bias, and a more uniform misclassification to either the neighboring fronto-parallel or exclusively in-depth motion stimulus **(Fig. 4A)**. To test for this bias, we computed the percentage of times oblique motion trials were misclassified as either the neighboring fronto-parallel or in-depth motion stimulus **(Fig. 4B)** based on the confusion matrices from the previous analysis **(Fig. 3C)**. We fit a linear mixed-effects model including the fixed effects of ROI (V1/hMT), motion direction (fronto-parallel/motion-in-depth), and their interaction. We also included by-subject random effects of intercept and slope. Misclassification percentage was the dependent variable. The model revealed a significant interaction between ROI (V1/hMT) and motion direction (fronto-parallel/motion-in-depth) on misclassification (*F*(1,1148) = 6.6192, *p* = .0102). A test of the simple effects showed a greater tendency in V1 to misclassify an oblique direction into its nearest fronto-parallel direction (*M*_*V1_fronto-parallel*_ = 24.61%, *SE*_*V1_fronto-parallel*_ = .84%) instead of its nearest motion-in-depth direction (*M*_*V1_motion-in-depth*_ = 18.36%, *SE*_*V1_ motion-in-depth*_= .87%; *F*(1,574) = 9.8682, *p* = .0018). Misclassification in hMT on the other hand was not significantly biased toward either direction (*M*_*hMT_fronto-parallel*_ = 20.36%, *SE*_*hMT_fronto-parallel*_ = .84%; *M*_*hMT_motion-in-depth*_ = 18.36%, *SE*_*hMT_ motion-in-depth*_= .92%; *F*(1,574) = .4629, *p* = .4965; **Fig. 4A**). The misclassification result suggests that V1 predominantly represents retinal motion signals, as would be expected from previous literature (Hubel and Wiesel 1974; Snowden et al. 1991; Wuerger, Shapley, and Rubin 1996). For hMT, the pattern of BOLD responses to oblique motion appears to be less determined by its fronto-parallel component, as would be expected for regions that contain more binocular neurons. The 3D trajectory is represented beyond simply encoding 3D motion based on the motion’s weaker fronto-parallel component. To further test this hypothesis, we conducted additional experiments that compared decoding performance to horizontal and vertical motion stimuli in V1 and hMT.

**Figure 4.**
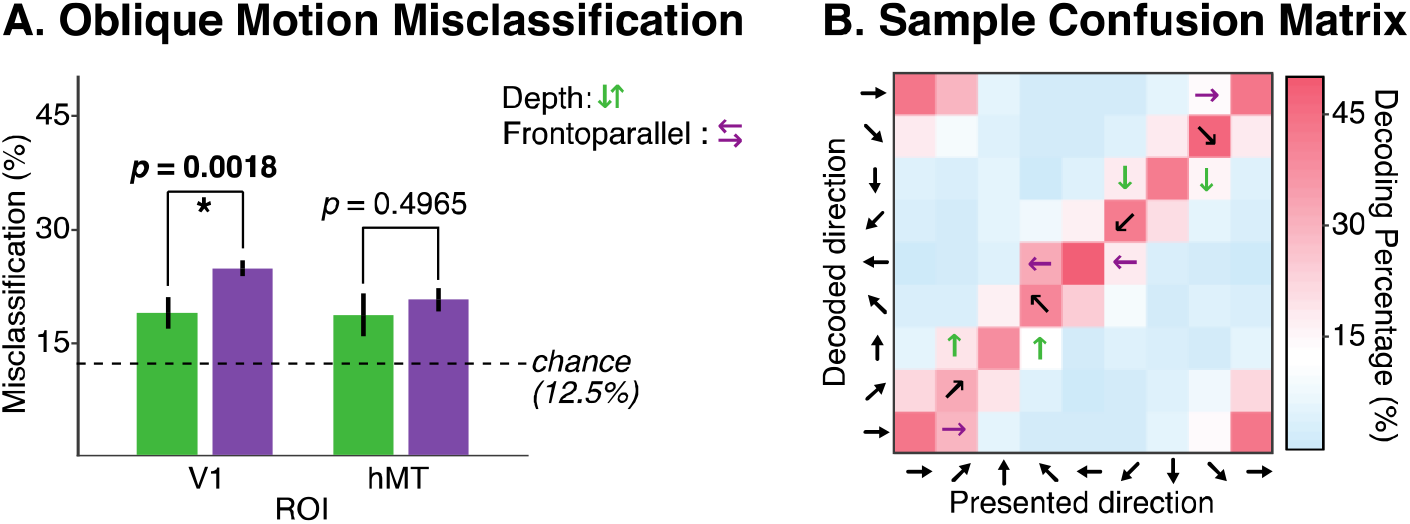
Decoding misclassification: Differences in decoding bias reveal differences in representation in V1 and hMT. ***A***, Oblique motion misclassification. Results are taken directly from the confusion matrices in Fig 3C. Green and purple bar each represent the percentage of the oblique trials that were misclassified as either the oblique motion’s depth or frontoparallel component. Dashed line: chance performance (12.5%). Error bars: SEM across subjects (*n*=9). ***B***, Sample confusion matrix. The results from the matrices that were included are indicated with purple (frontoparallel) and green (depth) arrows.

### Decoding horizontal versus vertical motion

Since our two eyes are horizontally offset, opposite retinal motion (leftward motion in one eye and rightward in the other) results from objects moving toward/away from the participant. In comparison, vertically opposite retinal motion (upward motion in one eye and downward in the other) is inconsistent with the world motion of a single stimulus. To dissociate the representation of retinal and world motion in cortical areas we therefore presented vertical rather than horizontal retinal motion to all participants in a control experiment. The vertical motion stimuli contained identical retinal motion energy, but produced percepts of transparent 2D rather than 3D motion. In ROIs that represent retinal motion, we expected decoding performance to be identical in the horizontal and vertical motion conditions. Conversely, in ROIs that represent world rather than retinal motion, we expected significantly better decoding performance in the horizontal compared to the vertical conditions.

The same nine subjects completed the same number of runs of the vertical condition and we decoded the vertical data using the same analysis pipeline. We fit a linear mixed-effects model including the fixed effects of ROI (V1/hMT), motion condition (horizontal/vertical), and their interaction. We also included by-subject random effects of intercept and slope. Decoding accuracy was the dependent variable. The model revealed a significant interaction between ROI (V1/hMT) and motion condition (horizontal/vertical) on decoding accuracy (*F*(1,1148) = 15.689, *p* < .0001). In V1, testing for simple effects showed decoding accuracy was not significantly different between horizontal (*M*_*V1_horizontal*_ = 29.64%, *SE*_*V1_horizontal*_ = .75%) and vertical motion conditions (*M*_*V1_vertical*_ = 26.97%, *SE*_*V1_vertical*_ = .76%; *F*(1,574) = 1.0071, *p* = .3011). In hMT on the other hand, we found significantly greater decoding accuracy in the horizontal motion condition (*M*_*hMT_horizontal*_ = 31.84%, *SE*_*hMT_horizontal*_ = .76%) compared to the vertical motion condition (*M*_*hMT_vertical*_ = 24.09%, *SE*_*hMT_vertical*_ = .65%; *F*(1,574) = 10.343, *p* = .0014; **Fig. 5**). Next, we asked if the pattern in hMT was driven by a specific motion type. We divided the eight motion directions into three groups: fronto-parallel, oblique, motion-in-depth. We tested if the difference between horizontal and vertical conditions in hMT was driven by one of these motion types. We fit a linear mixed-effects model of motion condition (horizontal/vertical), motion group (fronto-parallel /oblique/ motion-in-depth), and their interaction with random effect of subjects. The model again revealed greater decoding performance in the horizontal versus the vertical condition (*F*(2,1722) = 11.142, *p* = .0009). However we found no evidence for an interaction between motion condition and motion group (*F*(2,1722) = 2.5735, *p* = .0766). This result suggests that the difference between conditions in hMT is consistent across all three motion types and was not driven by any particular type. The results are consistent with the misclassification biases reported in the previous section. Taken together, these findings suggest that the signals in V1 predominantly represent retinal motion while the signals in hMT at least in part represent world motion.

**Figure 5.**
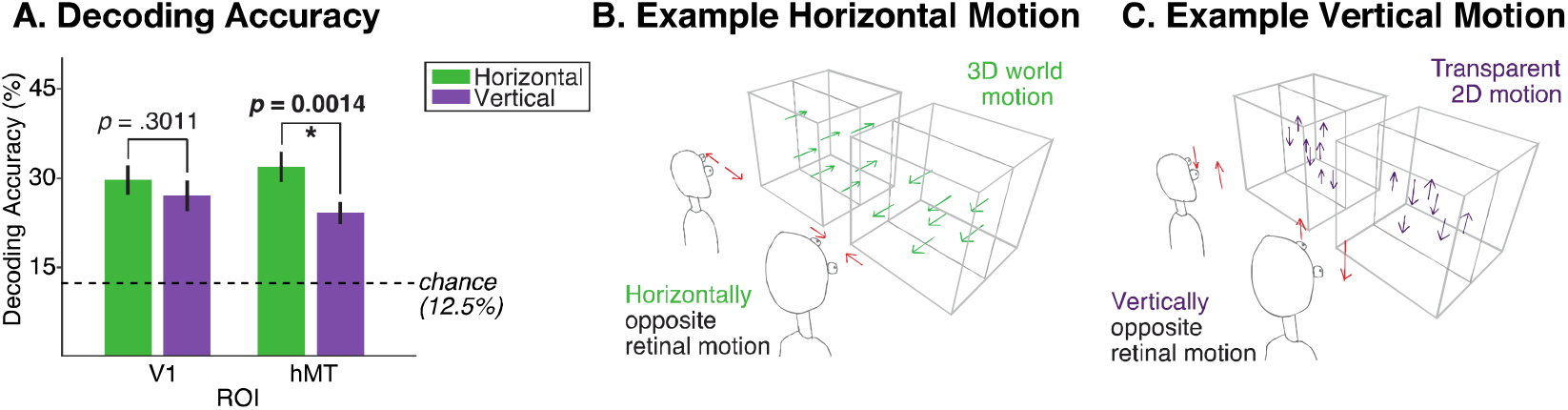
***A***, Decoding accuracy in the horizontal and vertical conditions. Dashed line: chance performance (12.5%). Error bars: SEM across subjects (*n*=9). ***B-C***, Example horizontal and vertical motion directions compared in this analysis.

### Motion decoding performance beyond V1 and MT

V1 and hMT are two critical regions for the processing of visual motion. However, the complete motion pathway involves additional areas. The classical V1 -> MT motion-processing pathway is based on studies of 2D motion in both humans and non-human primates. Our current work aims to advance our understanding of the motion pathway that underlies 3D motion perception. Thus, in the next sections, we look beyond V1 and hMT. We first investigate the role of ROIs in retinotopically-characterized visual cortex. Subsequently, we search for evidence for the representation of 3D motion signals beyond those ROIs in a searchlight analysis across the entire cortical surface.

### Representation of motion signals in retinotopically-characterized cortex

We decoded motion signals in 20 additional ROIs using our analysis pipeline (**Fig. 6A)**. Note that the V1 and hMT results in **Fig. 6A** are identical to those in **Fig. 5**. Many regions showed above-chance decoding accuracy in both the horizontal and vertical conditions, which was not expected for some regions, especially those located in the ventral pathway. Decoding accuracy also varied across the cortex. Unsurprisingly, a one-way ANOVA revealed a significant difference in decoding accuracy across ROIs for both conditions (*F*_*horizontal*_(21,176)= 6.77, *p* < .0001; *F*_*vertical*_(21,176)= 5.59, *p* < .0001). Early visual areas as well as dorsal regions tend to have relatively greater decoding accuracy compared to the ventral regions in the horizontal condition. To identify regions that show greater representation of world motion over retinal motion, we then ran a paired sample *t*-test of the horizontal decoding accuracy against the vertical conditions. In addition to hMT, three ROIs showed a significant difference between the horizontal and vertical conditions. We found significant differences between conditions in V3, hMT, V3A, and IPS0 (*t*_*V3*_(8) = 3.02, *p* = .0166; *t*_*hMT*_(8) = 3.03, *p*= .0163; *t*_*V3A*_(8) = 3.53, *p* = .0077; *t*_*IPS0*_(8) = 3.19, *p* = .0127), corrected for FDR using the Benjamini-Hochberg procedure (*q* = 0.1). These regions are colored in yellow in **Fig. 6B** and they are clearly not randomly distributed across visual cortex.

**Figure 6.**
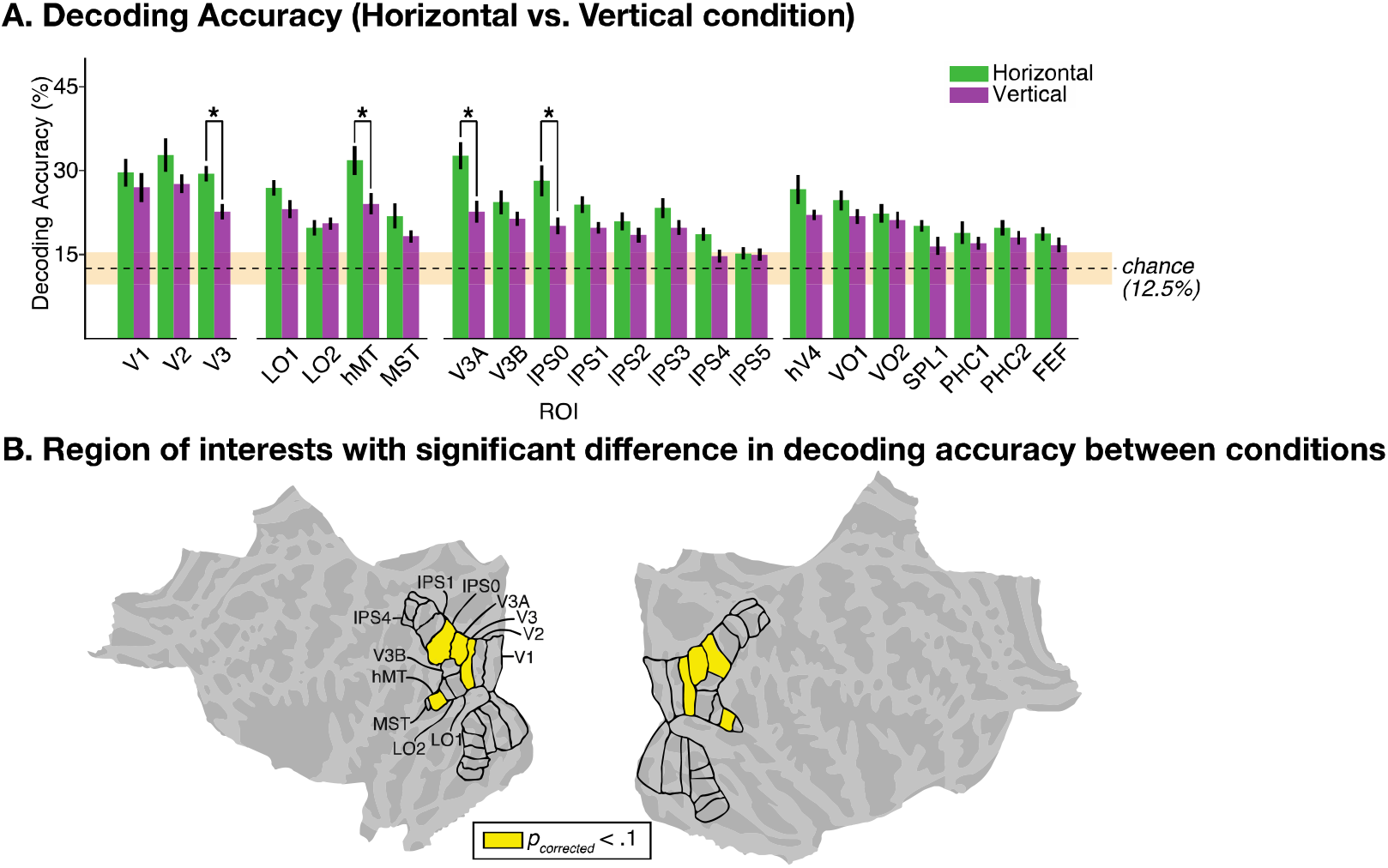
Motion decoding accuracy between the horizontal and vertical conditions. ***A***, Decoding accuracy averaged across subjects. Significant paired-sample t-tests between horizontal and vertical conditions are indicated at the top of the bars (* : *p*_*corrected*_ < .1; *df* = 8). Dashed black line: chance performance (12.5%). Light orange horizontal patch: 95% confidence interval of chance (9.55-15.25%). Error bars indicate SEM across subjects (*n*=9). ***B***, Regions of interest with significant differences in decoding accuracy between conditions colored in yellow in both hemispheres. The black outlines indicate each of the 22 ROIs.

As we expected early visual cortex to encode retinal motion, we found that decoding accuracy for vertical and horizontal motion did not differ significantly in V1 and V2 (*t*_*V1*_(8) = 0.95, *p* = .3718; *t*_*V2*_(8) = 1.94, *p*= .0879). We also found no significant differences in ventral regions such as PHC1-2, which are not commonly thought to be motion responsive. The differences between conditions start to arise in later dorsal regions. V3, V3A, and IPS0 are adjacent to one another and could process similar cues in our stimulus.

The preceding analyses were restricted to ROIs within the retinotopically mapped visual processing hierarchy. However, hMT appears relatively isolated, and one might have expected a role for MST as well. However, MST contains significantly fewer than 200 voxels (32 voxels on average) and this might have obscured results. Additionally, the cortical areas neighboring hMT are not typically identified using retinotopic mapping, but may nonetheless show evidence for the representation of motion signals. We therefore carried out a searchlight analysis to identify additional regions in which retinal and/or world motion signals may be reliably decoded.

### Searchlight classification

Our analyses so far have focused on visual ROIs that are part of the well-established parieto-occipital and temporo-occipital hierarchy and have previously been shown to contain clear retinotopic organization. To investigate the representation of binocular motion signals in additional visual, or indeed any cortical areas, we conducted a searchlight analysis across the entire cortical surface. We used a MATLAB classifier (see Materials and Methods) in the searchlight analysis instead of TAFKAP due to the latter’s high computational demand. This approach also verifies that our results do not depend on a particular choice of multivariate pattern analysis algorithm. Cross-validated classifier decoding results averaged across subjects are shown in **Fig. 7**. Every vertex served as the center of a searchlight sample consisting of the nearest 100 vertices. Inspecting the entire cortical surface, the accuracy ranged from 9.6% to 24.1% (*M* = 14.5% and *SD* = 1.41%) in the horizontal condition and from 10.31% to 23.58% (*M* = 14.8% and *SD* = 1.72%) in the vertical condition. The decoding accuracy of each sample was then compared with the bootstrapped chance distribution to calculate the *p*-value, which was then represented at each vertex. Despite variability across subjects, most subjects displayed the same pattern: reliable decoding performance around the canonical motion processing pathway (V1-hMT) as well as IPS0. The whole pathway is evident in the averaged map for the horizontal condition (**Fig. 7A**). Large patches cover the entirety of the early visual areas, the dorsal pathway, and regions around the hMT complex.

**Figure 7.**
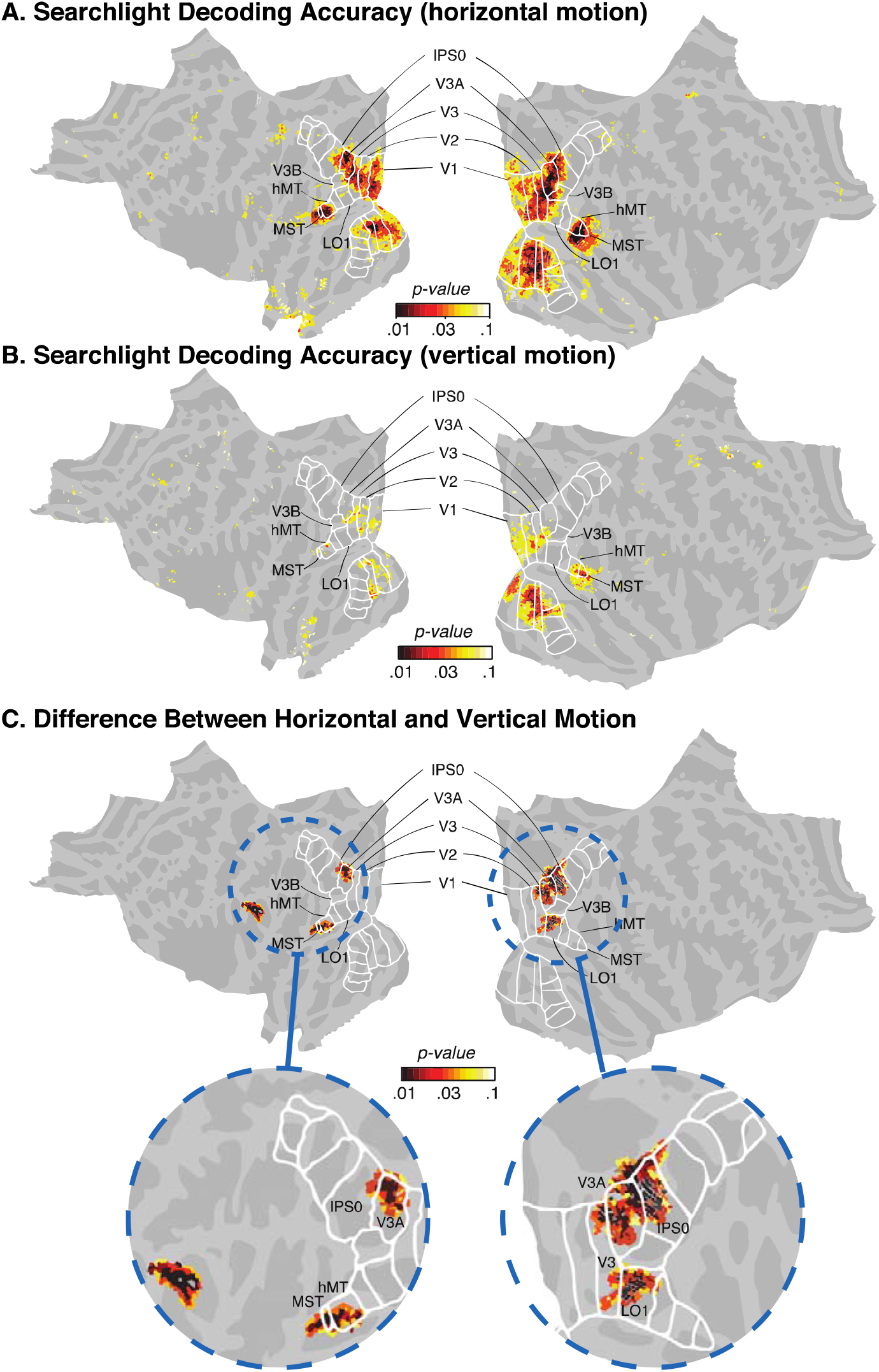
Searchlight decoding results on the cortical surface. ***A***, Classification accuracy for the horizontal motion condition. ***B***, Classification accuracy for the vertical motion condition. ***C***, The difference in decoding accuracy between the horizontal- and vertical-motion conditions thresholded by cluster size. The color bar indicates *p*-values in log scale and ranges from log(0.01) to log(0.1) for all three panels. The white outlines indicate each of the 22 ROIs. The blue dashed circles zoom into the significant clusters on the cortical surface.

To highlight regions that represent binocular motion signals beyond their retinal constituents, we contrasted decoding performance between the horizontal and vertical conditions.

We bootstrapped the data 5000 times and defined the *p*-value as the proportion of times the randomized condition difference was larger than the observed difference. For every random sample, we found the largest significant cluster across both hemispheres and recorded the size of that cluster into our reference distribution of significant cluster sizes. After 5000 resamples, we found that the largest cluster size resulting from random sampling was 66 vertices. We thresholded our data using this criterion, leading to the map of *p*-value results in **Fig. 7C**. This result revealed five major clusters. Laterally on the left hemisphere, we identified a region anterior to hMT extending anteriorly into the posterior part of the middle and superior temporal sulcus and ending at the supramarginal gyrus. We did not see the same pattern in the right hemisphere. Instead, we identified a region in LO1. Two additional clusters are both located around V3A and IPS0, with the regions in the left hemisphere centering around V3A and the right hemisphere centering around both V3A and IPS0. The cluster around V3A/IPS0 is the largest cluster identified by the searchlight result. Interestingly, no early visual area (V1-3) showed up on this map, consistent with the rest of our results (except for V3) and suggesting that these regions decode vertical motion equally well as horizontal motion. This also indicates that the significant difference between the conditions in V3 (see **Fig. 6**) might arise from signals in V3A instead.

### Motion direction preferences

Previous research has suggested that high decoding accuracy of neural signals in early visual areas might arise from confounds such as global preference maps for both orientation and motion (Beckett et al. 2012; Wang et al. 2014). Thus, we used additional analyses to scrutinize the representation of motion signals in V1 and hMT that could underlie the decoding results. The motion-preference map (**Fig. 8**) revealed a horizontal fovea-inward bias pattern along the majority of the visual pathway except in peripheral early visual areas (V1-3), where a horizontal fovea-outward bias was observed instead. Horizontal fovea-inward biases are indicated by a pattern where vertices consistently shows the largest response to motion towards the fovea. We observe that the left hemisphere (i.e., the right visual field) prefers leftward motion (motion towards the fovea), and the right hemisphere (i.e., the left visual field) prefers rightward motion in foveal parts of V1, V2, and V3, as well as V3AB, LO1-2, hMT, MST, and IPS0. The peripheral regions of V1, V2, and V3 show a fovea-outward bias where each vertex prefers motion away from the fovea. In addition to the predefined ROIs, an area anterior to IPS5 and an area anterior and inferior to hMT and MST also showed horizontal fovea-inward bias. Previous work showed that retinotopically biased preference contributes to the decoding of stimulus orientation (Freeman et al. 2011) and motion direction (Beckett et al. 2012; Wang et al. 2014). We see that the stimulus location only affected bias early in the visual hierarchy, which explains the high decoding accuracy in V1 for the horizontal motion condition.

**Figure 8.**
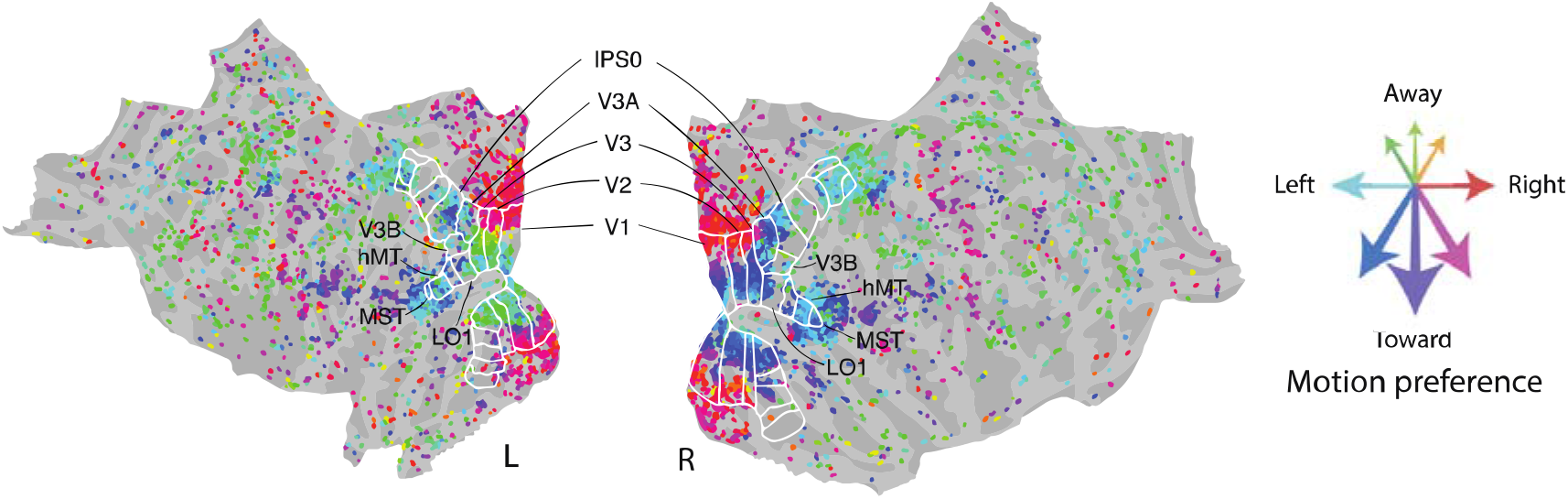
Motion direction preferences averaged across nine subjects on a flattened map for both hemispheres. The color of each dot represents the preferred motion direction for that specific vertex after averaging across subjects. The color bar ranges from red (rightward motion) to green (away motion) to teal (left motion) and to purple (toward motion) in a counter-clockwise manner on the HSV colormap. Black contours indicate the outline of ROIs. The results are thresholded by only showing vertices in the top tenth percentile of averaged and z-scored BOLD responses.

## Discussion

We presented independent motion signals to the two eyes to identify the cortical regions that underlie the transformation from retinal to world motion signals. We identified three major clusters. In early visual cortex (V1-V3), we found no significant difference in decoding performance between horizontal (3D) and vertical (transparent) motion directions, suggesting a representation of retinal motion. In voxels in and surrounding hMT however, decoding performance was consistently superior for horizontally opposite compared to vertically opposite motion stimuli. We also identified a third cluster in V3A/IPS0 that showed a similar pattern. Our results reveal the parts of the visual processing hierarchy that are critical for the transformation of retinal into world motion signals and suggest a role for V3A/IPS0 in the representation of 3D motion signals in addition to its sensitivity to 3D object structure and static depth.

### Representation of retinal motion signals in early visual cortex

We found that early visual areas V1-3, and especially V1, yield a strong representation of retinal motion and very little sign of the representation of 3D world motion. The stimuli in the vertical and horizontal condition contained the same retinal motion energy. Although vertical motion does not produce a coherent world-motion percept, in these regions we were able to decode vertical transparent motion just as well as 3D horizontal motion.

Additionally, we found a strong bias towards decoding 2D motion in V1. When oblique motion directions are misclassified, they are more likely to be classified based on their 2D retinal motion signals. For example, the response in V1 to rightward-away motion more closely resembles rightward than away motion. This makes sense if a region is at a stage of the visual processing hierarchy where the motion signals from the two eyes have yet to merge into a unified percept. Thus, our results suggest that early visual areas encode retinal motion signals, consistent with prior work on component motion signals (Britten et al. 1993; Movshon and Newsome 1996; Born and Bradley 2005; Rust et al. 2006).

### Representation of motion signals in later regions

Our hMT results suggest a mixture of retinal and world-motion representations. The eight-way decoding result shows that hMT can decode horizontal 3D motion well, but the performance dropped significantly when decoding vertical 2D motion. hMT also did not show a bias towards 2D motion in the misclassification results, similar to IPS0.

It is important to consider what information is represented in hMT. Motion processing begins in V1 with 2D direction selectivity as well as tuning for other properties (size/spatial frequency, disparity, etc.). Downstream, higher-level motion cues must be derived, such as interocular velocity difference (IOVD) and changing disparity over time (CD). These cues are then integrated to allow the brain to reconstruct the world around us. Motion direction can be reliably decoded throughout many of these motion-processing stages. It is important to distinguish exactly what information is exploited by the decoder. hMT showed a significantly different pattern of results compared to V1 in our study. We made a clear distinction that V1 encodes retinal motion signals and hMT does not. However, we cannot say if hMT fully represents world motion from these results alone. The same argument holds for the other regions (IPS0, V3A, and LO1) that are highlighted in our studies. These regions showed a different pattern of results compared to early visual areas, but additional data are needed to elucidate the form of the representation of motion signals in these areas.

Area LO1 is located between V3A and hMT on the lateral side of the cortex, which overlaps with the kinetic occipital region (KO; (Tyler et al. 2006; Larsson and Heeger 2006). The literature suggests that area KO is sensitive to both motion and depth cue-defined kinetic boundaries (Van Oostende et al. 1997; Zeki, Perry, and Bartels 2003). Although we have no stimulus-defined boundaries or contours in our stimuli, it is possible that we were decoding the stereoscopic cues that LO1 uses for boundary perception. KO is also thought to define boundaries from discontinuities in motion direction. Our stimulus changes motion direction regularly and abruptly. If there were a pause between different intervals of each motion direction, perhaps this region would not be activated as prominently. It is also worth noting that this region does not respond uniquely to motion – shapes defined by color contrast and motion contrast result in equal sensitivity (Van Oostende et al. 1997; Zeki, Perry, and Bartels 2003).

Our findings of significant decoding of 3D motion signals using stereoscopic visual cues in IPS0 were somewhat surprising, especially in the absence of evidence for the representation of retinal motion signals. No previous research has suggested that IPS0 is part of the motion-processing pathway. Instead, IPS0 is a highly multisensory area and frequently appears in studies of working memory (Bray et al. 2015; Brigadoi et al. 2017), motor planning (Buneo and Andersen 2006), spatial representation (Makin, Holmes, and Zohary 2007), and multimodal integration (Regenbogen et al. 2018; Ladda, Wallwork, and Lotze 2020). Future work is needed to determine if the areas involvement is purely visual, or contains a representation of 3D space across multiple different modalities, such as 3D motion auditory cues.

The remaining question circles back to potential differences in the representations in IPS0 and the classical motion region hMT. Representation of binocular cues in hMT is critical for many visual tasks other than motion perception, such as structure from motion and depth perception (Bradley, Chang, and Andersen 1998; Orban et al. 1999; Backus et al. 2001; Neri, Bridge, and Heeger 2004; Preston et al. 2008). These cues become meaningful after the integration across the two retinas and can be used by multiple other regions such as MST, LO1, V3A, and IPS0.

Could the decoder have exploited alternative sources of information? One possibility to consider is binocular disparity, particularly because our horizontal motion stimuli contain (horizontal) binocular disparities, whereas our vertical motion stimuli do not. It is therefore a reasonable concern that the observed differences in decoding performance between horizontal and vertical motion condition were due to binocular disparity rather than motion cues, especially in regions like V3A and hMT, that are known to be sensitive to static binocular disparity signals. However, these binocular disparity signals by themselves are insufficient to discriminate between the presented motion directions. For example, rightward motion and leftward motion stimuli contained identical disparity cues, but were nonetheless discriminable by our decoder. Regions that only encode static depth and not world motion would not produce superior decoding accuracy in the horizontal condition.

A second potential contributor to differences in decoding accuracy between conditions is the arrangement of monocular and binocular neurons in different visual areas. Regions containing predominantly monocular neurons will have less conflicting signal within each neuron. In the most extreme case, half of the neurons in this region receive input from the left eye and the other half receives input from the right eye, then MVPA could exploit this eye-specific information to make classification of the world motion direction even if this region never forms any coherent world-motion percept. A difference in the proportion of monocular neurons between regions could result in the difference in decoding performance. However, this contributor is most impactful in single-neuron studies. With fMRI, the voxel is the smallest unit of signals, and each voxel contains millions of neurons. It would be much more difficult for the MVPA classifier to exploit the arrangement of monocular neurons at the voxel level. On the other hand, if our classifier was indeed exploiting eye-specific motion signals, then we would not see a difference in decoding performance between the horizontal and vertical motion conditions as the motion energy contained within each eye was identical.

In conclusion, multivariate pattern analysis methods have been highly successful in decoding sensory input based on neural signals. However, it has not always been clear in previous work exactly what signals any particular decoder exploits. In the domain of orientation and motion specifically previous work has shown that artifacts (vignetting) may drive decoding performance. Here we provide one way to dig deeper. By carefully manipulating visual stimuli and evaluating the impact on decoder performance across conditions, we can start to identify the representational transformations in cortex that must underlie behavior. We hope to extend this method to other stimulus properties and modalities and move beyond the reconstruction of sensory input from neural signals and towards identifying the representational transformations that underlie perceptual inference.

## Acknowledgements

Support: Global PhD Student Fellowship to PW. NIH EY08266 to MSL. We thank Ari Rosenberg and Lowell Thompson for valuable comments on the manuscript.

## Footnotes

1 See supplementary video 1 for horizontal stimuli movie: https://www.dropbox.com/s/apeoz0X4ixp5mry/Horizontal%20Condition.mp4?dl=0.

2 See supplementary video 2 for vertical stimuli movie: https://www.dropbox.com/s/i5t5ovbrcozezer/Vertical%20Condition.mp4?dl=0.

## Notes

### Competing Interest Statement

The authors have declared no competing interest.

